# Intrinsic water use efficiency depends on stomatal aperture rather than stomatal density in C_3_ and C_4_ grasses grown at glacial CO_2_ and low light

**DOI:** 10.1101/2021.07.22.453465

**Authors:** Walter Krystler Israel, Alexander Watson-Lazowski, Zhong-Hua Chen, Oula Ghannoum

## Abstract

We investigated how stomatal morphology and physiology control intrinsic leaf water use efficiency (*iWUE*) in grasses. Two C_3_ and six C_4_ grasses were grown at ambient (400 µl L^-1^) or glacial CO_2_ (180 µl L^-1^) and high (1000 µmol m^-2^ s^-1^) or low light intensity (200 µmol m^-2^ s^-1^). C_4_ grasses tended to have higher *iWUE* and CO_2_ assimilation rates, and lower stomatal conductance (g_s_), operational stomatal aperture (*a_op_*) and guard cell K^+^ influx rate relative to C_3_ grasses, while stomatal size (SS) and stomatal density (SD) did not vary according to the photosynthetic type. Overall, *iWUE* and g_s_ depended most on *a_op_* and density of open stomata. In turn, *a_op_* correlated with K^+^ influx, stomatal opening speed on transition to high light and SS. Species with higher SD had smaller and faster-opening stomata. Although C_4_ grasses operated with lower g_s_ and *a_op_* at ambient CO_2_, they showed a greater potential to open stomata relative to maximal stomatal conductance (g_max_), indicating heightened stomatal sensitivity and control. We uncover novel links between *a_op_*, g_s_, *iWUE* and K^+^ influx amongst grasses and differential K^+^ influx responses of C_4_ guard cells to low light, revealing molecular targets for breeding crops with high *iWUE*.

**Highlights:** Across C_3_ and six C_4_ grasses, intrinsic water use efficiency was strongly associated with stomatal conductance, operational stomatal aperture, guard cell K^+^ influx and stomatal opening speed on transition to high light.

## INTRODUCTION

A key desirable feature of future ‘climate-ready’ crops is improved water use efficiency (WUE) to cope with decreased water availability due to a rising global population, shrinking arable lands, increasing temperature and declining rainfall (Flexas, 2016). Crop WUE is governed by multiple factors including leaf-level intrinsic WUE (*iWUE*) (Condon *et al*., 2004; Passioura, 1977). Leaf *iWUE* is defined as the ratio of CO_2_ assimilation rate (A_net_) over stomatal conductance (g_s_) to water vapour (Farquhar and Sharkey, 1982). Photosynthesis and g_s_ are closely correlated over the long term (Wong *et al*., 1979; Pinto *et al*., 2016) such that selecting for reduced g_s_, as a means of improving *iWUE*, often leads to reduced A_net_ and productivity (Blum, 2009; Ghannoum, 2016). However, g_s_ and A_net_ can be uncoupled in the short term and under various conditions (McAusland *et al*., 2016; von Caemmerer *et al*., 2004) raising the prospect of improving stomatal traits without affecting CO_2_ fixation, and hence yield (Ghannoum, 2016; Webster *et al*., 2016). Therefore, developing the next generation of smart crops requires a greater understanding of how stomata regulate *iWUE*. Plant breeders often utilise variations within a narrow set of crop-related species to identify improved traits. The downfall of this strategy is the limited pool of genetic diversity available in a single species. Mining natural variation among diverse grasses will increase the potential of identifying beneficial stomatal traits (Anderson *et al*., 2016; Cano *et al*., 2019; Reeves *et al*., 2018; Watson-Lazowski *et al*., 2019).

By controlling CO_2_ entry into the leaf, stomata provide the initial limitation of CO_2_ uptake. Grasses possess unique stomata composed of two dumbbell-shaped guard cells flanking subsidiary cells (Nunes *et al*., 2020; Rudall *et al*., 2017). Stomatal opening and closing govern gas exchange and are regulated by the movement of solutes and water in and out of the guard cells. These sophisticated structural and functional features enable fast and tight stomatal control in response to environmental fluctuations (Cai *et al*., 2017; Chen and Blatt, 2010; Chen *et al*., 2010; Franks and Farquhar, 2007). They also contribute to the success, productivity and survival of grasses in semi-arid and arid habitats (Chen *et al*., 2017; Taylor *et al*., 2018; Taylor *et al*., 2012). Therefore, it is not surprising that grasses including some of the most important cereal crops have been domesticated by our ancestors. Many grasses, including major crops (e.g., maize, sugarcane, sorghum and switchgrass) assimilate CO_2_ using the C_4_ photosynthetic pathway (Brown, 1999; Sage, 2004). C_4_ crops are becoming increasingly important for food and bioenergy security, with the global production of C_4_ maize currently surpassing that of key C_3_ cereals such as wheat and rice (Varshney *et al*., 2012).

C_4_ photosynthesis operates a carbon concentrating mechanism (CCM) which elevates CO_2_ concentration ([CO_2_]) around the carbon-fixing enzyme Rubisco (ribulose 1,5-carboxylase/oxygenase) in the bundle sheath, thereby minimizing photorespiration and allowing higher photosynthetic rates at relatively low stomatal conductance (Hatch, 1987; Long, 1999). Consequently, *iWUE* is generally higher in C_4_ relative to C_3_ grasses (Taylor *et al*., 2012). C_4_ CCMs are broadly classified according to the major C_4_ decarboxylase utilised, either solely or with a secondary decarboxylase (Kanai and Edwards, 1999; Watson-Lazowski *et al*., 2018; Wingler *et al*., 1999); nicotinamide adenine dinucleotide-malic enzyme (NAD-ME), nicotinamide adenine dinucleotide phosphate-malic enzyme (NADP-ME), and phosphoenol carboxykinase (PCK). Apart from differences in leaf anatomy and biochemistry, the three C_4_ subtypes also show variations in geographic distribution according to rainfall and resource (e.g. light, water, nitrogen) use efficiency (Ghannoum *et al*., 2011; Ghannoum *et al*., 2005; Hattersley, 1992). In particular, NAD-ME grasses exhibit higher *iWUE* at glacial [CO_2_] (gCO_2_: 180 µl l^-1^) (Pinto *et al*., 2014) and higher plant WUE under water stress (Ghannoum *et al*., 2002) relative to NADP-ME and PCK counterparts. Recently, we identified two subtype-specific differentially expressed transcripts linked to stomatal function. βCA1, conserved only in NAD-ME species, is a gene known to sense CO_2_ in plants; and SASP, conserved only in PCK species, is involved in controlling stomatal aperture (Watson-Lazowski *et al*., 2020). Hence, the first objective of this study was to elucidate how stomatal control of *iWUE* differs among grasses depending on the photosynthetic pathway (C_3_ versus C_4_) or the C_4_ biochemical subtypes.

Light regulates stomatal function via photosynthesis-dependent and independent processes (Assmann and Jegla, 2016; Lawson, 2009). The photosynthesis-mediated (or red light) response affects g_s_ by driving photosynthetic linear electron transport and is partially mediated by decreasing intercellular [CO_2_] (C_i_) (Baroli *et al*., 2008; Messinger *et al*., 2006; Mott, 2009). In addition, guard cells directly respond to red□light, independent of mesophyll photosynthesis, though the signal has not been conclusively identified (Busch, 2014; Lawson and Matthews, 2020). Blue light directly affects the phototropins in guard cells through photosynthesis-independent reactions leading to the activation of plasma membrane H^+^-ATPase pumps (Christie, 2007; Hiyama *et al*., 2017; Shimazaki *et al*., 2007). The hyperpolarised or depolarised plasma membrane activates inward- or outward-rectifying K^+^ channels, which are the key regulators of guard cell homeostasis, leading to stomatal opening or closure, respectively (Hosy *et al*., 2003; Kim *et al*., 2010; Schroeder and Fang, 1991). Photosynthesis and stomatal conductance respond with different speed to short-term changes in light intensity, causing transient disequilibrium in *iWUE* (Lawson and Vialet-Chabrand, 2019; McAusland *et al*., 2016). Hence, the rates of stomatal closure or opening in response to light transients are valuable traits for optimised *iWUE* (Deans *et al*., 2019; Knapp and Smith, 1987). Following long-term exposure to high light, stomatal density (SD) and stomatal pore lengths increase in certain species, such as tomato (Gay and Hurd, 1975), tobacco (Thomas *et al*., 2003), *Arabidopsis ssd-1* mutants (Schluter *et al*., 2003) and *Eucalytus globulus* (James and Bell, 2000). Conversely, decreased SD and stomatal index (SI), but not stomatal size (SS) were observed in tobacco leaves (Gerardin *et al*., 2018; Thomas *et al*., 2003) and *Coffea arabica* (Pompelli et al., 2010) exposed to shade. However, it remains unclear if and how C_3_ and C_4_ stomata respond differently to light. Accordingly, our second objective was to determine how stomatal responses to short-term light and long-term light changes differ among C_3_ and C_4_ grasses, including the three C_4_ subtypes.

A period of low (glacial) atmospheric [CO_2_] (gCO_2_) during the Oligocene is thought to be the primary driver for the evolution of C_4_ photosynthesis (Christin *et al*., 2008; Sage, 2004). Plants adapted to the geological decline in atmospheric [CO_2_] by having higher SD compared to current atmospheric [CO_2_] (Woodward, 1987; Woodward and Kelly, 1995). In response to short-term (instantaneous) changes in [CO_2_], stomata respond by opening with decreasing [CO_2_] and by closing with increasing [CO_2_] both in darkness and light (Assmann, 1999; Azoulay-Shemer *et al*., 2016; Lawson *et al*., 2011). In the long term, stomata acclimate to changes in [CO_2_] through biochemical and anatomical changes which may modify the instantaneous responses. Growth at gCO_2_ increased g_s_ in C_3_ and C_4_ grasses (Maherali *et al*., 2002; Pinto *et al*., 2014; Sage and Coleman, 2001), and increased SD by 42% in *Arabidopsis* relative to ambient CO_2_ (Li *et al*., 2014). Conversely, growth at elevated [CO_2_] reduced SD, g_s_, and sensitivity to [CO_2_] (Ainsworth and Rogers, 2007; Leakey, 2009; Maherali *et al*., 2002; Woodward and Kelly, 1995; Xu *et al*., 2016). To date, few studies investigated the mechanisms underpinning the acclimation of C_3_ and C_4_ stomata to [CO_2_]. Hence, our third objective was to determine how stomatal responses to short-term light and long-term light changes to [CO_2_] differ among C_3_ and C_4_ grasses, including the three C_4_ subtypes.

To address these objectives, we grew closely related C_3_ and C_4_ grasses with different biochemical subtypes under gCO_2_ (180 CO_2_ µl L^-1^) and/or low light (200 µmol m^-2^ s^-1^) as well as control conditions (1000 µmol m^-2^ s^-1^ of light and 400 µl CO_2_ L^-1^) with the aim of further exploring stomatal responses to these environmental stimuli. Light is expected to impart a greater stomatal and photosynthetic response than CO_2_ because plants are continuously exposed to light fluctuations in their open environment, while atmospheric CO_2_ gradually changed over time. Nevertheless, stomata have been shown to respond to both factors in the short and long term. Hence, we hypothesised that stomata will respond to gCO_2_ by increased aperture and density (*Hypothesis 1*) while stomata respond to low light by reduced aperture and density (*Hypothesis 2*). We also hypothesised that both low light and gCO_2_ will affect transient stomatal responses (*Hypothesis 3*). Elucidating these hypotheses will shed light on how stomatal traits (morphology, physiology, dynamics and/or ion flux) control *iWUE* in C_3_ and C_4_ grasses acclimated to low light and gCO_2_. In turn, such new understanding will be useful for developing future cereal crops with improved *iWUE*.

## MATERIALS AND METHODS

### Plant materials and grow conditions

Eight C_3_ and C_4_ grasses (Poaceae) representing different photosynthetic and biochemical subtypes were grown in walk-in growth chambers (Biochambers, Winnipeg, Manitoba) controlled by LI-820 CO_2_ gas analysers (LI-COR Inc., Lincoln, NE, USA) as described in (Watson-Lazowski *et al*., 2020). They were two C_3_: *Panicum bisulcatum* (Thunb.) and *Steinchisma laxa* (Zuloaga); two C_4_ NAD-ME: *Panicum miliaceum* (L.) and *Leptochloa fusca* [(L.) Kunth.]; two C_4_ NADP-ME: *Panicum antidotale* (Retz.) and *Setaria viridis* [(L.) Beauv.]; and two C_4_ PCK: *Megathyrsus maximus* (Jacq., synonym: *Panicum maximum*) and *Chloris gayana* (Kunth.). Two species, *L. fusca* (NAD-ME) and *C. gayana* (PCK) belong to subfamily Chloridoideae while the remaining six species belong to Subfamily Panicoideae (Grass Phylogeny Working, 2012). All species were individually planted in 3 L pots containing soil (Osmocote^®^ Professional Seed Raising Mix, Scotts, Australia) and trace elements supplement (Osmocote^®^ Plus Trace Elements, Scotts, Australia). Seeds were germinated in the control chamber (400 µl CO_2_ L^-1^ and 1000 µmol m^-2^ s^-1^). After germination, plants were transferred to respective growth chambers with different light intensity (high: 1000 µmol m^-2^ s^-1^ or low: 200 µmol m^-2^ s^-1^ at pot level) and CO_2_ concentration (ambient: 400 µl l^-1^ or glacial: 180 µl L^-1^) treatments. High and low light treatments were abbreviated as HL and LL, respectively. Ambient and glacial [CO_2_] were abbreviated as aCO_2_ and gCO_2_), respectively. The average day/night temperature was 28/22^○^C, relative humidity (RH) was 60%, and the photoperiod was 14 h. Plants were watered to full capacity every 1-2 days, fertilised as required and randomized twice a week to reduce the effects of within-chamber variation. There were 3-5 biological replicates per treatment.

### Leaf gas exchange

Five weeks after germination, gas exchange was measured with LI-6400XT infrared gas analyser (LI-COR Inc., Lincoln, NE, USA) using the youngest fully expanded leaf from the main stem. Cuvette conditions were maintained at 60-65% RH, 28^○^C leaf temperature, and 350 mol s^-1^ flow rate. Two types of measurements were taken between 10:00 and 15:00 inside the growth chambers for all eight grasses. Steady-state measurements of net CO_2_ assimilation rate (A_net_), stomatal conductance (g_s_), intercellular CO_2_ (C_i_) and intrinsic water use efficiency (*iWUE* = A_net_/g_s_) were made at the four respective growth conditions. Measurements were also made at light-saturated (2000 µmol m^-2^ s^-1^) and low CO_2_ (180 µl L^-1^ CO_2_) conditions to induce maximal stomatal conductance (g_sat_) and light-saturated photosynthesis (A_sat_). Prior to each measurement, leaves were pre-adapted at the chamber conditions for 10-15 min until steady-state rates of CO_2_ and H_2_O exchange were reached. At least eight technical replicates across 3-5 biological replicates per species were measured.

### Stomatal morphology using epidermal impressions

Stomatal impressions were characterised based on the methods described by (Sekiya and Yano, 2008; Weyers and Johansen, 1985) with some modifications. Epidermal imprints containing stomatal patterns were taken using clear nail varnish from the adaxial and abaxial surfaces of all eight grass species using the same leaf where the steady-state gas exchange was measured. Dried nail varnish impressions were transferred to glass slides (Knittel Glass, Germany), and images were captured using Nikon microscope equipped with CCD camera and DS-U3 controller (NIS-F1 Nikon, Tokyo, Japan). Images were captured under a field of view of 0.0768 mm^2^ at 200× and 400× total magnification and processed using Nikon NIS

Element imaging software (Nikon, Tokyo, Japan). To ensure uniformity, stomatal images were captured between two major veins of the leaf lamina and were analysed using Image J software (NIH, USA). Ten anatomical stomatal parameters (Caine *et al*., 2019; Gerardin *et al*., 2018) were measured (Figures S1 and S2): guard cell width (GCW), guard cell length (GCL), subsidiary cell width (SCW), aperture width (AW), aperture length (AL), operational stomatal aperture (*a_op_*), stomatal size (SS), stomatal density (SD), stomatal index (SI), and density of open stomata (OD). Anatomical SS and *a_op_* were calculated by assuming the area of an ellipse whereby:

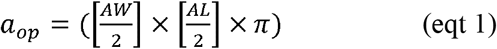

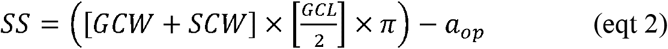

such that AW/2 and GCW+SCW were the short axes while AL/2 and GCL/2 were the long axes of *a_op_* and *SS,* respectively. SD was calculated as the number of stomata per unit area while OD was calculated by counting the number of open stomata (i.e., showing a gap between the two guard cells) per unit area as observed from the nail varnish impressions (Figure S2). A stoma was counted if more than 50% was located within the field of view. SI was calculated as the ratio of stomata to total cell numbers in the epidermis (stomata plus non-stomatal cells) × 100. Three random microscopic fields from the adaxial and abaxial surfaces per biological replicate were measured totalling >300 stomata per species in each growth chamber and the mean of the adaxial and abaxial surfaces were used (*leaf side p = 0.75, df = 1,7*).

### Maximal stomatal conductance using epidermal peels

Maximum stomatal aperture (*a_max_*) was measured following (Liu *et al*., 2014; Mak *et al*., 2014) with some modifications. The mid-portion of the second fully expanded leaf was harvested and soaked immediately in stomata opening buffer (50 mM KCl, 5 mM Na-MES, pH 6.1) after which the leaf epidermis was promptly peeled. Peels were mounted in a 35-mm petri dish with 0.13 mm glass bottom (MatTek Corp, MA, USA) coated with silicone adhesive (B-521, Factor II, Lakeside, AZ 85929), and then 2 ml of opening buffer was added. Epidermal peels were incubated for 2 hr at 700 µmol m^-2^ s^-1^ irradiance to induce maximal stomatal opening*. a_max_* was calculated using the same equation (eqt 1) for calculating *a_op_*. Maximum stomatal conductance (*g_max_*) was calculated using *a_max_* determined from epidermal peels. One-sided *g_max_* was calculated using the equation presented by (Franks and Beerling, 2009a) based on the (Brown and Escombe, 1901) diffusion model of gases for plants:

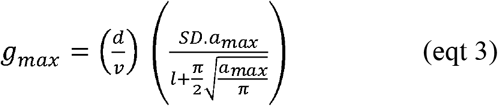

where *d* = diffusivity of water vapor at 25^○^C (0.0000249 m^-2^ s^-1^); *v* = molar volume of air (0.0224 m^3^ mol^-1^); *SD* = stomatal density (m^-2^); *a_max_* = maximum stomata aperture (m^2^) and *l* is the stomata pore depth which is assumed to be equivalent to guard cell width (m). The total, two-sided *g_max_* was calculated as *g_max_* (abaxial) × 2 since both leaf sides had similar SD.

### Stomatal responses to light transitions

The rates of stomatal closure in response to transition to LL (100 µmol m^-2^ s^-1^) followed by rate of stomatal opening in response to transition to HL (1,000 µmol m^-2^ s^-1^) were measured under similar conditions as steady-state measurements for four Panicoidea species: *P. bisulcatum* (C_3_), *P. miliaceum* (C_4_-NAD-ME), *P. antidotale* (C_4_-NADP-ME) and *M. maximus* (C_4_-PCK). Initially, leaves were pre-adapted at 400 µl CO_2_ l^-1^ and 1000 µmol quanta m^-2^ s^-1^ for 20-30 min until reaching steady-state CO_2_ uptake. Subsequently, steady-state g_s_ was auto-logged every 10 s for 15 min after which light intensity was reduced to 100 µmol m^-2^ s^-1^. The exponential decay of g_s_ was monitored every 10 s until a new steady-state was reached (∼20 min). Light intensity was then increased to 1000 µmol m^-2^ s^-1^ to monitor the rate of stomatal opening (Figure S3). The rate of stomatal closure was fitted using an exponential decay model as described by (Elliott-Kingston *et al*., 2016):

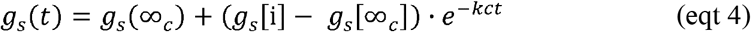

where *g_s_*(*t*) is the stomatal conductance at time point (*t*), *g_s_* (∞_c_) is the stomatal conductance at the new steady-state after irradiance was switched to 100 µmol m^-2^ s^-1^, *g_s_*[i] is the initial steady-state conductance at *t* = 0, *kc* is the exponential decay constant for closing, and the closing half-time (c*t_1/2_*) is the time required for *g_s_* to decrease by 50% of the difference between its initial and final values. The rate of g_s_ opening was fitted using an exponential rise to maximum model:

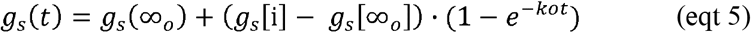

where *g_s_*(*t*) is the stomatal conductance at time point (*t*), *g_s_* (∞_o_) is the stomatal conductance at the new steady-state after irradiance was switched back to 1000 µmol m^-2^ s^-1^, *g_s_*[i] is the initial steady-state conductance at 100 µmol m^-2^ s^-1^, *ko* is the exponential rise constant for opening, and opening half-time (o*t_1/2_*) is the time required for *g_s_* to increase by 50% of the difference between its initial and final values. Lower *t_1/2_* indicates a faster response time. Closing and opening half times (*t_1/2_*) were calculated using the respective exponential decay or rise constants (*k*) as:

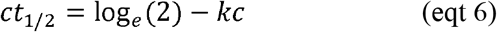

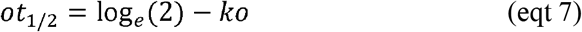

### Estimation of water and CO_2_ losses during light transitions

Excess transpiration (ΔW, Figure S4a), the transpiration above final steady-state transpiration due to slow stomatal closure during transitions from HL (1000 µmol m^-2^ s^-1^) to LL (100 µmol m^-2^ s^-1^) was estimated according to (Deans *et al*., 2019). It was defined as the time-integrated difference between the initial and final steady-state transpiration rates (*E*):

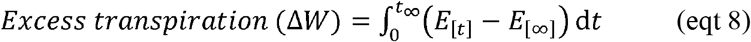

where *E*[t] was the initial steady-state *E* at 1000 µmol m^-2^ s^-1^ and *E*[∞] was the transpiration rate at time point 90% (t_90_) wherein 90% of the residual *E* is no longer changing under the new steady-state (100 µmol m^-2^ s^-1^). Potential forgone potential photosynthesis (ΔC, Figure S4b) due to slow photosynthetic induction during transitions from LL to HL was defined as the time-integrated difference between the final and initial A_net_ rates according to (Deans *et al*., 2019):

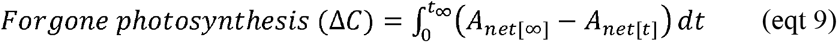

where A_net_[∞] was the final steady-state assimilation rate at HL (1000 µmol m^-2^ s^-1^) and A_net_[t] was the residual steady-state assimilation rate at LL (100 µmol m^-2^ s^-1^).

### Microelectrode ion flux estimation

Fluxes of K^+^, a major ion responsible for stomatal control, were measured for four Panicoidea species: *P. bisulcatum* (C_3_), *P. miliaceum* (C_4_-NAD-ME), *P. antidotale* (C_4_-NADP-ME) and *M. maximus* (C_4_-PCK). Measurements were made using microelectrode ion flux estimation on guard cells of leaf epidermal peels (Zhao *et al*., 2019) with some modifications. Prior to measurements, ion-selective microelectrodes were prepared as described by (Chen *et al*., 2005) and subsequently backfilled with a solution of 200 mM KCl. The tip was broken to achieve a 2-3 µm diameter and was front-filled with potassium ionophore I (Sigma, Switzerland), a liquid ion exchanger to achieve high resistance (4–6 GΩ). The microelectrodes were then mounted into the micromanipulator holder with AgCl wire together with a reference microelectrode backfilled with 1 M KCl in 2% agar. The microelectrodes were calibrated using 2, 5, and 10 mM KCl achieving a calibration slope of - 55 mV and a correlation coefficient ≥0.999.

Leaf epidermal peels were soaked in OB and promptly peeled to ensure stomatal integrity. The peels were mounted on a glass cover slide using a silicon prosthetic adhesive (Dow-Corning, USA) and immersed in a chamber filled with the stomatal measuring solution (10 mM KCl, 0.5 mM CaCl_2_, 5 mM MES-KOH, pH 6.1) in a 45-degree position. The peels equilibrated in the measuring solution were focused on the same plane together with the microelectrodes that were focused on top of the guard cells and moving in two positions, near the cells (10 µm) and away from the cell surface (40 µm) in 5-sec cycle, 80 µm amplitude for 10 min. The steady-state ion flux was calculated using the basic planar diffusion geometry described by (Newman, 2001) via the MIFEFLUX software and expressed per epidermal peel surface area. The same leaf and species used for g_s_ kinetics measurements were assayed.

### Statistical analyses

ANOVA was performed using linear mixed-effects models (*lme*) in R (V.3.4.2; R Core Team, 2017). The full factorial experimental design included light (2 treatments) CO_2_ (2 treatments) and photosynthetic type or subtype, where species were nested inside the photosynthetic type (2 groups, n = 2 C_3_ and n = 6 C_4_ species) or biochemical subtypes (3 groups, n = 2 C_4_ species per subtype). There were 3-5 plants (biological replicates) per species, and at least 3 technical replicates per plant (unbalanced ANOVA). Homoscedasticities and normalities were checked by examining the quantile plots and the re-fitted when necessary. Since the comparisons consisted of hierarchal (nested) terms, the *F*-statistic was re-calculated such that the signal (numerator mean squares) to noise (denominator mean squares) ratio reflected the true inference on the effects of [CO_2_], light intensity, type/subtype and their respective interactions. The effect of glacial [CO_2_] treatment was the average effects of chambers with gCO_2_ regardless of light conditions (HL+gCO_2_ and LL+gCO_2_) while the effect of ambient [CO_2_] was the average effects of aCO_2_ chambers (HL+aCO_2_ and LL+aCO_2_) regardless of light treatment. Low light effect was the average effect of the chambers with LL conditions (LL+aCO_2_ and LL+gCO_2_) while high light effect was the mean variation within chambers with HL conditions (HL+aCO_2_ and HL+gCO_2_). The equivalent *p value* was calculated for a given degree of freedom. When significant, means were ranked using Tukey’s *post-hoc* at a stringency of α=0.05. Linear correlations between two variables were expressed in *r^2^* followed by significance values. Figures were plotted using the *ggplot2* package (Wickham, 2016). Principle components analysis (PCA) was generated in R using the *pca.comp* function, scaling each variable using the scale=TRUE function, then using *ggbiplot* to look for relationships between samples and traits.

## RESULTS

### Glacial CO_2_ and low light differentially affect leaf gas exchange of C_3_ and C_4_ grasses

When measured under growth conditions, photosynthetic rate (A_net_) and stomatal conductance (g_s_) showed significant [CO_2_], light and photosynthetic type, but not subtype, effects (Table 1). In the six C_4_ grasses, A_net_ was higher under HL and reduced to a greater extent by LL (-63%) relative to the two C_3_ (-44%) species (Figure 1a, Tables 1 and S1). In all species, g_s_ was greater under gCO_2_ relative to aCO_2_, while g_s_ was lower under LL relative to HL only in C_4_ species, which maintained lower g_s_ relative to C_3_ counterparts under gCO_2_ and/or LL (Figure 1b, Tables 1 and S1). Intrinsic water use efficiency (*iWUE*) was higher in the C_4_ than C_3_ grasses, and lower under gCO_2_ relative to aCO_2_, but was not affected by LL (Figure 1c, Tables 1 and S1).

**Fig. 1.**
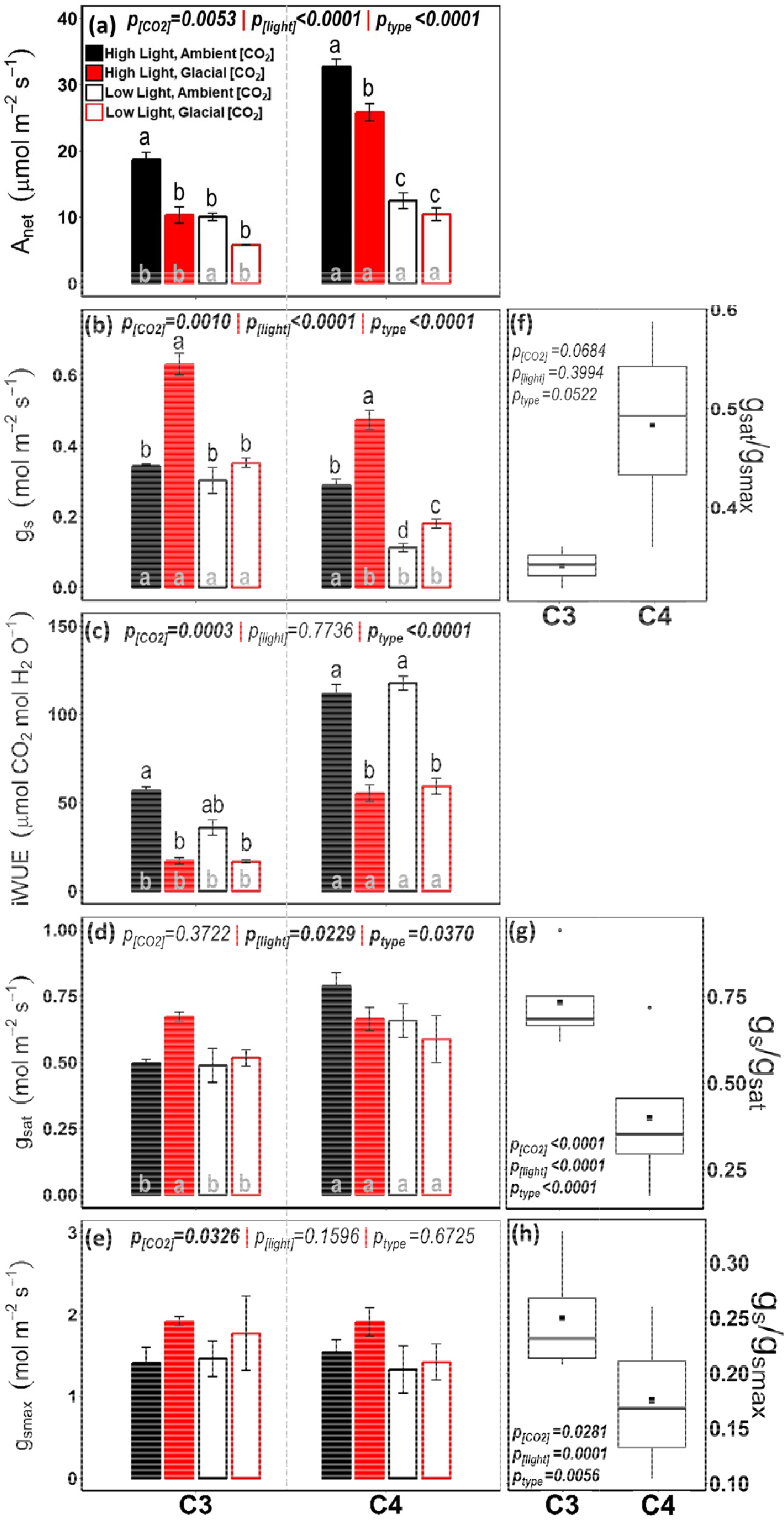
Effects of glacial [CO_2_] and/or low light on leaf gas exchange parameters of two C_3_ and six C_4_ grasses (two of each subtype): (**a**) Net CO_2_ assimilation rate at growth conditions (A_net_), (**b**) stomatal conductance at growth conditions (g_s_), (**c**) maximum stomatal conductance calculated from anatomical traits (g_max_), (**d**) light-saturated stomatal conductance at gCO_2_ (g_sat_), and (**e**) intrinsic water-use efficiency (*iWUE* = A_net_/) measured at growth conditions. Data represent means ± SE (n = 2 C_3_ or 6 C_4_) with species as replicate within each group. Means with the same letter are not significantly different at *p* 0.05 using Tukey’s HSD *post hoc*. Letters on top of the columns describe treatment effect for each group. Letters inside the columns indicate group differences at each of the growth condition. Plots **f-h** show the distribution of (**f**) g_sat_/g_max_, (**g**) g_s_/g_sat_ and (**h**) g_s_/g_max_. The box and whiskers represent the 25-75 and 0-100 percentiles, respectively. The square and line inside the box represent the mean and median of the data, respectively.

**Table 1.**
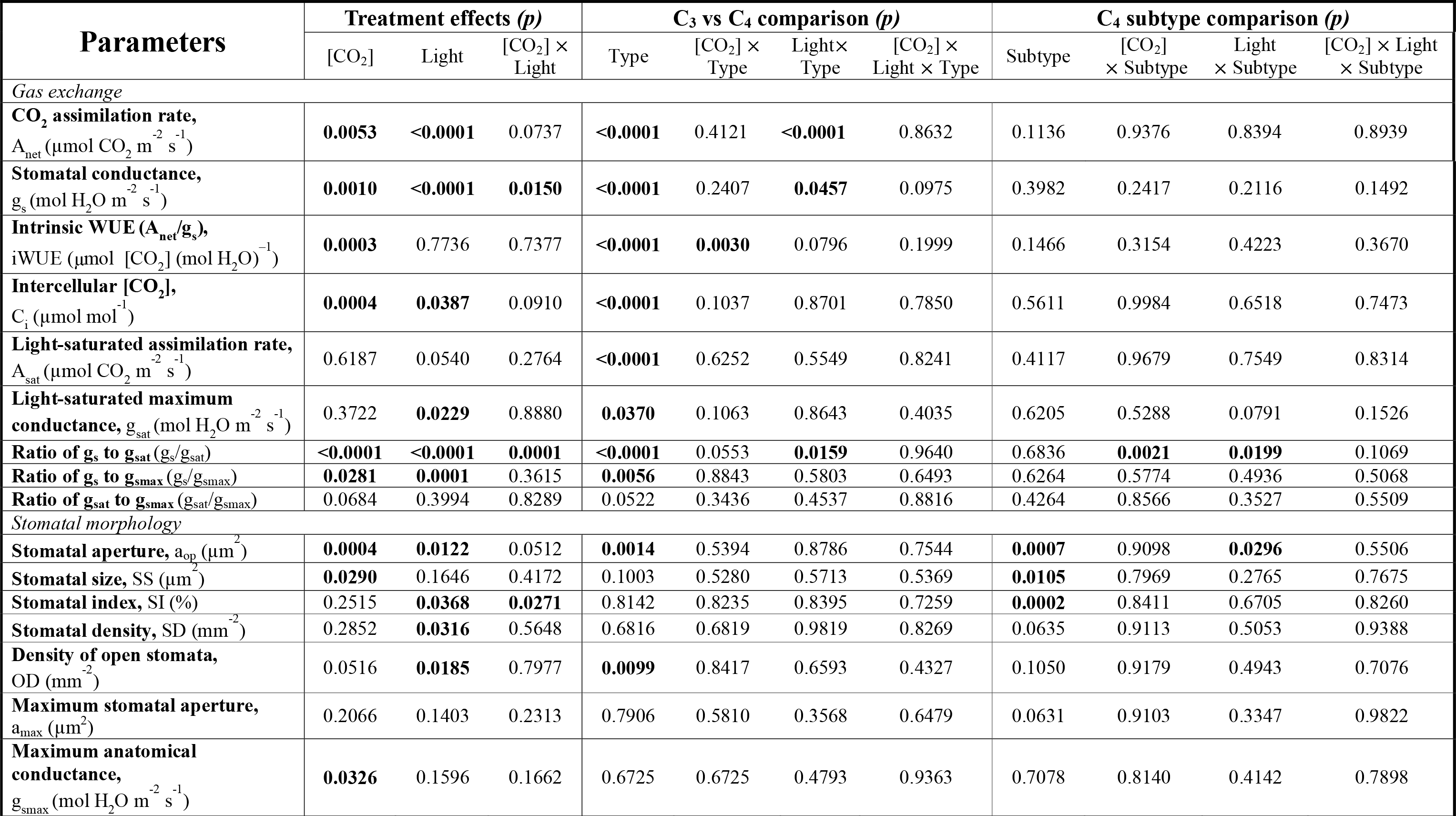
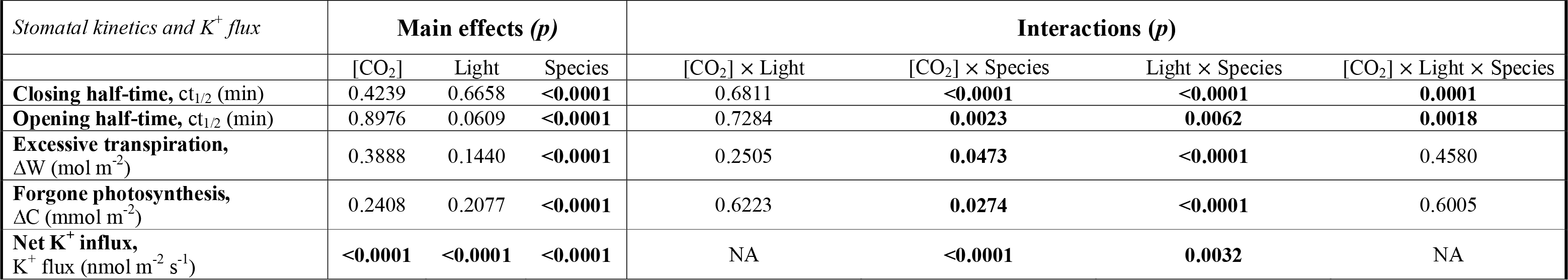
Statistical analysis summary of the main effects and interactions of CO_2_, light, species, and photosynthetic type or subtype. For leaf gas exchange and stomatal morphology parameters, two separate *glm* analyses were used with eight grass species nested in either Type (2 C_3_ and 6 C_4_) or Subtype (n=2 in each of the three subtypes). Species was used as a random effect while the functional type was utilised as the fixed effect. For stomatal kinetics and guard cell K^+^ influx parameters the analysis used species (n=4), [CO_2_] light. Bold numbers indicate significant effect at *p*≤0.05.

When measured under saturating light (2000 µmol m^-2^ s^-1^) and low [CO_2_] (180 µl L^-1^) to encourage stomatal opening, light-saturated photosynthesis (A_sat_) and stomatal conductance (g_sat_) were higher in C_4_ relative to C_3_ species, and tended to be lower under LL relative to HL in most C_4_ species. Both A_sat_ and g_sat_ showed no significant CO_2_ effect (Figure 1d, Tables 1 and S1). Under control conditions, the g_s_/g_sat_ ratio averaged 0.7 and 0.4 in C_3_ and C_4_ species, respectively (Figure 1g, Tables 1 and S1). The g_s_/g_sat_ ratio was higher under gCO_2_ in all species (reaching close to 1 in C_3_ species), and lower under LL in C_4_ species, relative to the control (Tables 1 and S1). These results indicate that C_4_ grasses operate with a much lower g_s_, relative to g_sat_, than C_3_ grasses (Figure 1g, Tables 1 and S1).

### Leaf surface morphological responses to glacial CO_2_ and low light were driven by stomatal aperture

The stomatal complexes of the grass species were amphistomatous and paracytic, composed of two lateral, non-oblique (parallel to guard cells) subsidiary cells (SC) (Figure S2). The operational stomatal aperture (*a_op_*) was the most responsive anatomically-measured traits in the grasses (Figure 2, Tables 1 and S2). Overall, *a_op_* and density of open stomata (OD) were slightly lower in C_4_ than C_3_ species, mostly due to lowest *a_op_* in the C_4_-PCK relative to the other species (Figure 2a,d, Tables 1 and S2). Within C_4_ species, SS was smaller in PCK and SI was higher in NADP-ME species (Figure 2, Tables 1 and S2). In partial support of *Hypothesis 1*, growth at gCO_2_ led to higher *a_op_* but had no effect on stomatal density (SD). Growth at gCO_2_ also led to higher maximal stomatal conductance (g_max_) and slightly lower stomatal size (SS) relative to the control treatment (Figures 1e and 2, Tables 1 and S2). In full support of our *Hypothesis 2*, growth at LL led to lower *a_op_*, SD, OD and stomatal index (SI) of most species (Figures 1e and 2, Tables 1 and S2).

**Fig. 2.**
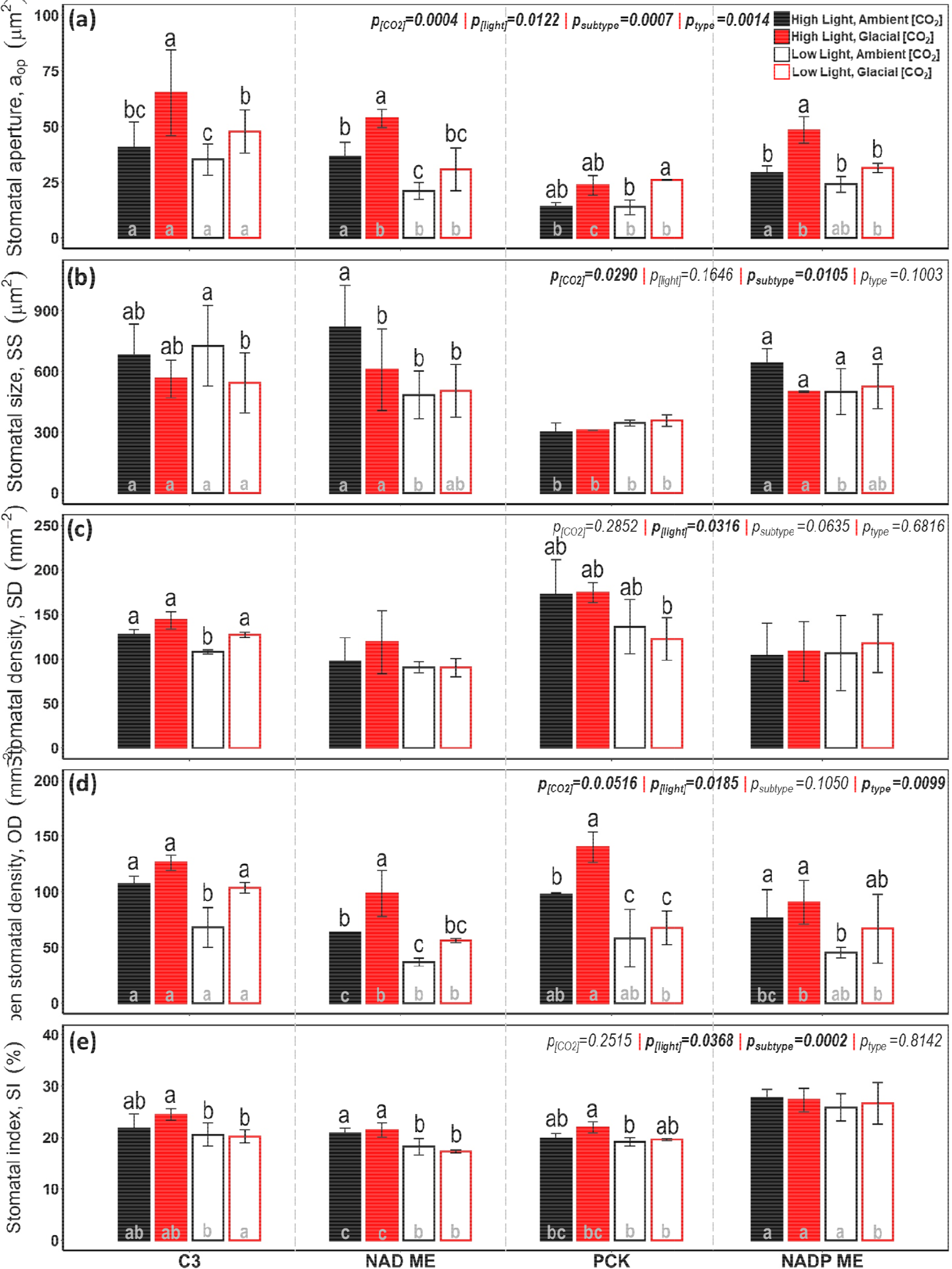
Acclimation of stomatal traits to glacial [CO_2_] and/or low light in two C_3_ and six C_4_ grasses (two of each subtype): (**a**) operational stomatal aperture (*a_op_*), (**b**) stomatal size (SS), stomatal density (SD), (**d**) density of open stomata (OD), (**e**) stomatal index (SI). Data represent means ± SE (n=2 species in each group). Means with the same letter are not significantly different at *p*≤0.05 using Tukey’s HSD *post hoc*. Letters on top of the columns describe treatment effect for each group. Letters inside the columns indicate group differences at each of the growth condition.

Physiologically induced maximal stomatal aperture (*a_max_*) was 3-5-fold greater than *a_op_* captured using *in situ* epidermal impressions (Table S2). Under control conditions, the g_s_/g_max_ ratio was slightly higher in C_3_ (0.24) than C_4_ (0.19) species (Figure 1h, Tables 1 and S1). The g_s_/g_max_ ratio was higher under gCO_2_ in all species, and lower under LL in the C_4_ species, relative to the control (Tables 1 and S1). The resultant g_sat_/g_max_ was relatively constant and lower in the C_3_ relative to the C_4_ species (Figure 1f, Tables 1 and S1). Hence, C_4_ grasses had a greater potential of stomatal opening (measured as g_sat_) relative to g_max_ (Figure 1f).

**Table 2.**
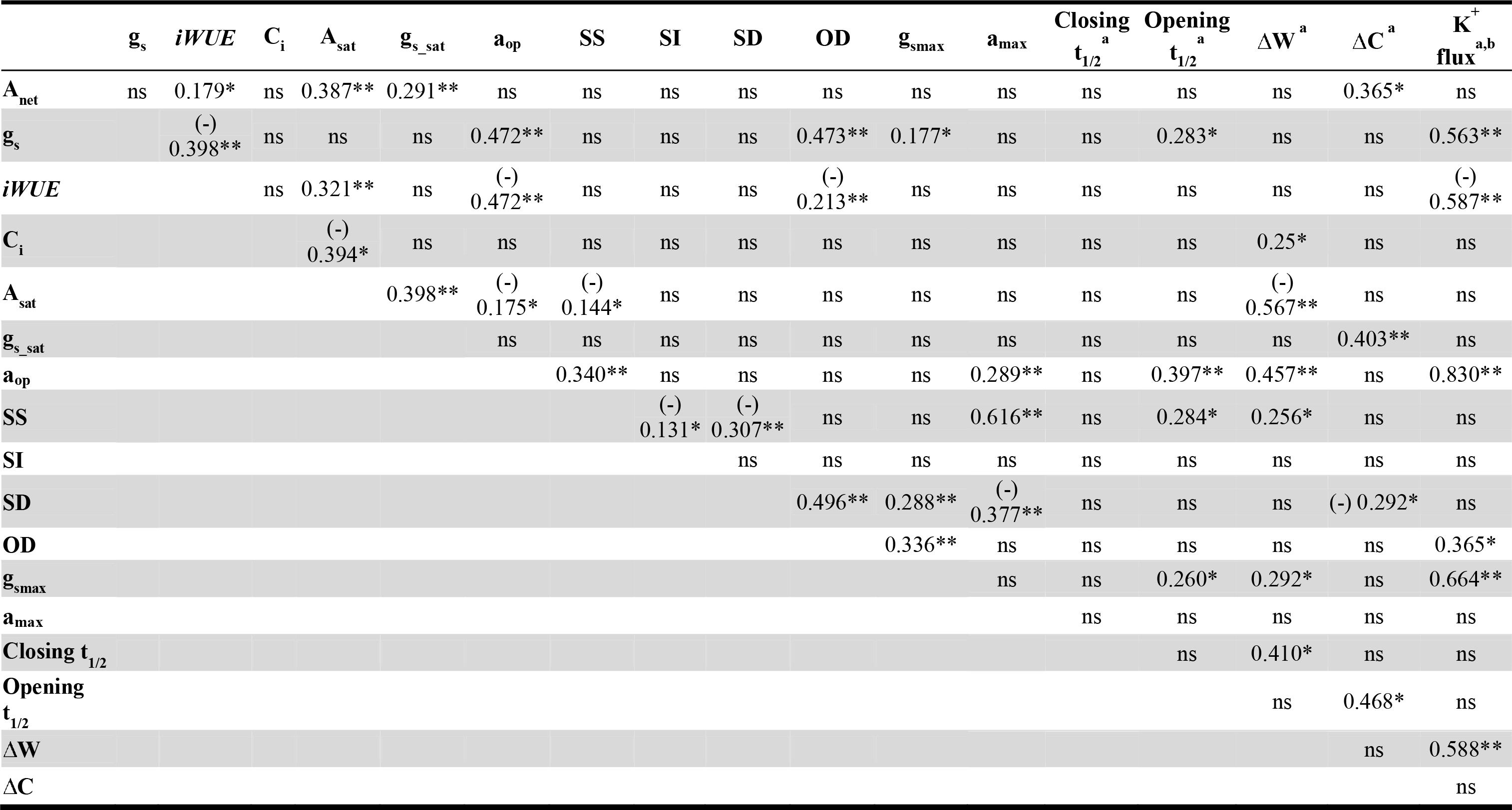
Correlation matrix between parameters among eight grasses acclimated to glacial [CO_2_] and/or low light. Linear regression direction is indicated in the parenthesis. Values represent r^2^ between two parameters. Significance codes are as follows: *significant at *p*≤*.0.05*; **significant at *p<0.01*; ^ns^not significant.^a^Only four *Panicoideae* species were analysed; ^b^Low light glacial CO_2_ treatment was not included.

### Correlations among stomatal variables of C_3_ and C_4_ grasses

Overall, g_s_ correlated best with *a_op_* (r^2^ = 0.472; Figure 3a) and OD (r^2^ = 0.473, Table 2), and weakly with g_max_ (r^2^ = 0.177, Table 2). In turn, *a_op_* correlated well with SS (r^2^ = 0.340; Figure 3b), and SS showed a negative trade off with SD (r^2^ = 0.307; Figure 3c). In addition, *a_max_* correlated positively with SS (r^2^ = 0.647) and *a_op_* (r^2^ = 0.378), and negatively with SD (r^2^ = 0.297) (Table 2). Maximal anatomical (g_max_) and diffusional (g_sat_) stomatal conductances did not correlate (Table 2). Moreover, g_max_ correlated best with OD (r^2^ = 0.336) due to a stronger correlation with SD (r^2^ = 0.288) than g_s_ (0.177) (Table 2). A good illustration of these correlations is within PCK species, which had smaller *a_op_* and SS, greater SD but similar g_max_ relative to other species (Figures 1 and 2, Tables 1 and S2).

**Fig. 3.**
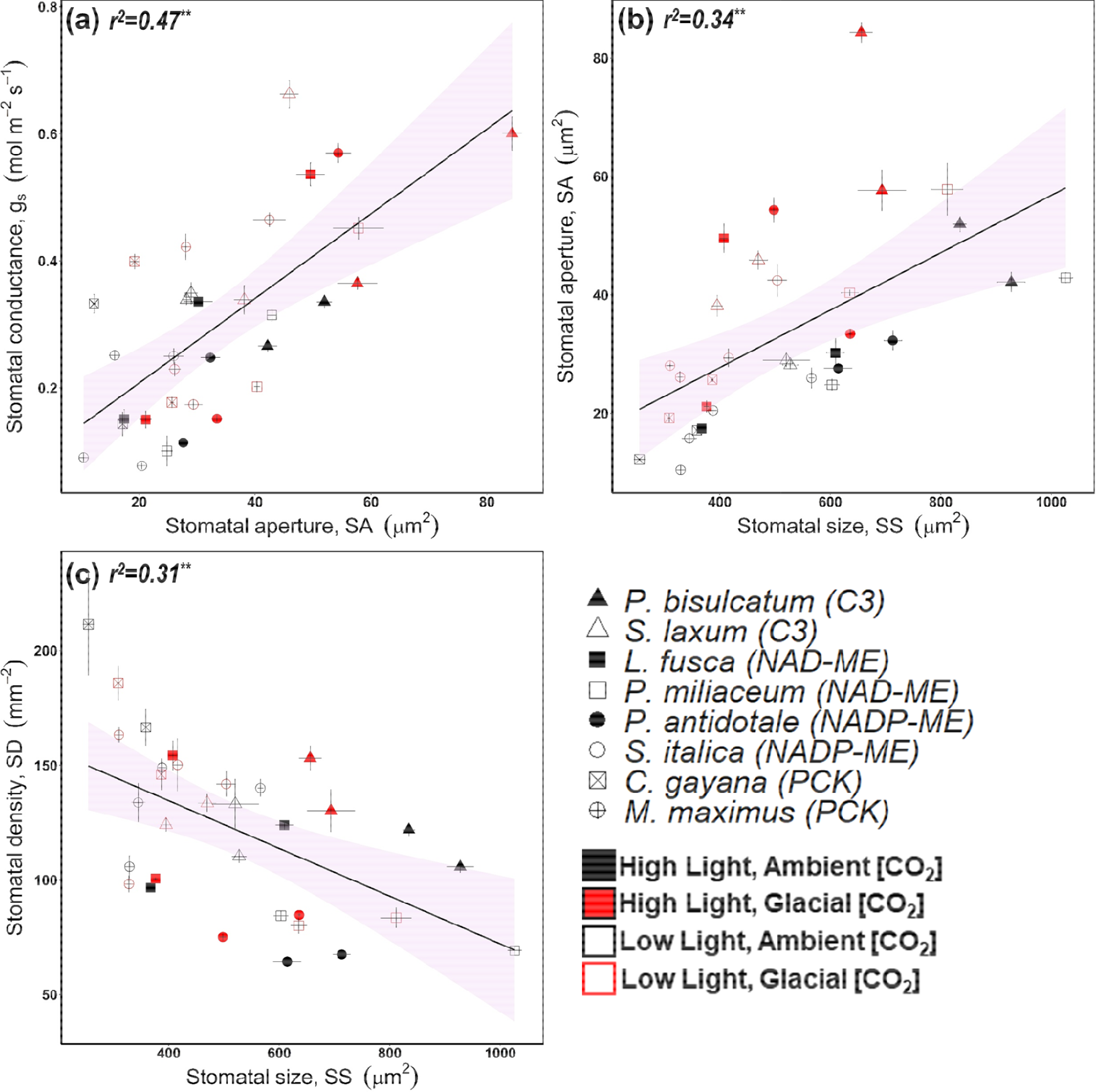
Relationships between various stomatal traits among two C_3_ and six C_4_ grasses (two of each subtype) acclimated to glacial [CO_2_] and/or low light. Points represent the species mean ± SE at each growth condition, which were indicated by different colours. Fitted linear models are as follows: (**a**) g_s_ = 0.0067 × *a_op_* + 0.0742; (**b**) *a_op_* = 0.0485 × *SS* + 8.3318; (**c**) *SD* = 176.19 - 0.1041 × *SS*. *\*p<0.05; **p<0.01; ***p<0.001*. Shaded regions represent the 95% confidence interval of the linear model using all the data.

### Smaller and more closed stomata opened faster in response to high light transition

Stomatal kinetics in response to short-term light transitions and guard cell K^+^ influx were measured in only one species per type or subtype (one C_3_ and three C_4_ grasses). Stomatal closing (ct_1/2_) and opening (ot_1/2_) half-times varied between grass species (Table 1). *M. maximus* had a lower ot_1/2_ relative to the other species (Figure 4a-b), consistent with its lower *a_op_* and SS (Figure 2a-b). In disagreement with our *Hypothesis 3*, ct_1/2_ showed no consistent response to growth conditions. In addition, ot_1/2_ was not affected by growth [CO_2_] and was marginally lower (*p* = 0.0609) under LL in only two C_4_ species relative to the control treatment (Figure 4a-b, Tables 1 and S3). ct_1/2_ and ot_1/2_ were not correlated, but ot_1/2_ positively correlated with *a_op_* (r^2^ = 0.378, Figure 4c) and to a lesser extent with SS (r^2^ = 0.284), g_s_ (r^2^ = 0.283) and g_max_ (r^2^ = 0.260), such that smaller and relatively closed stomata opened faster in response to increased light intensity during measurements (Table 2).

**Fig. 4.**
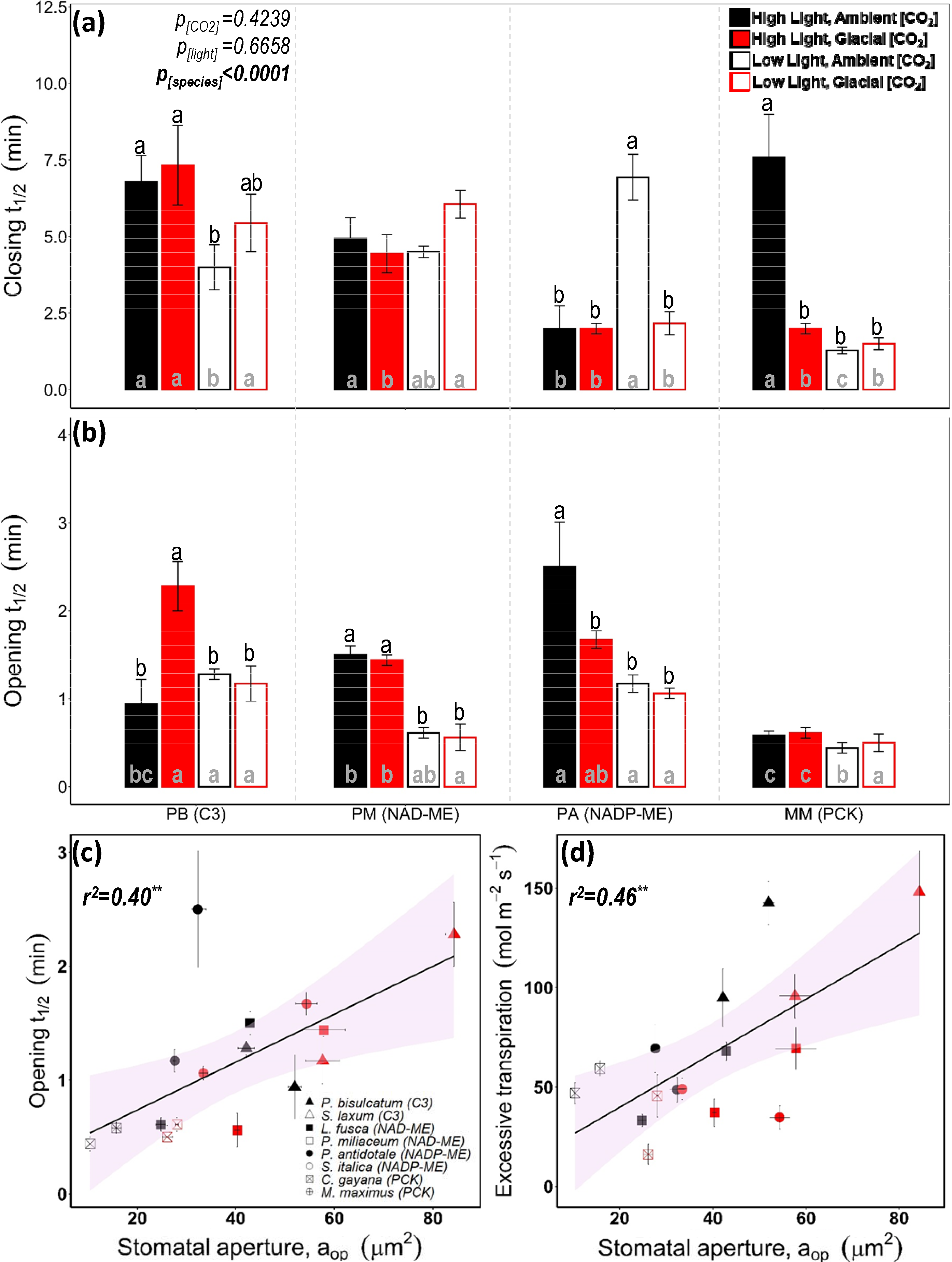
Stomatal (**a**) closing half-time and (**b**) opening half-time in response to light transitions for four grass species acclimated to glacial [CO_2_] and/or low light. Means with the same letter are not significantly different at *p*≤0.05 using Tukey’s HSD *post hoc*. Letters on top of the columns describe treatment effect for each species. Letters inside the columns indicate species differences at each of the growth condition. Species abbreviations: *P. bisulcatum* (PB, C_3_), *P. miliaceum* (PM, C_4_-NAD-ME), *P. antidotale* (PA, C_4_-NADP-ME) and *M. maximus* (MM, C_4_-PCK). Linear correlations (**c**) between opening half-time and stomatal aperture and **(d)** between excessive transpiration and stomatal aperture. Fitted linear models are as follows: (**c**) *Ot_1/2_* = 0.021 × *a_op_* + 0.3165; (**d**) Δ*W* = 1.3594 × *a_op_* + 12.634. *\*p<0.05; **p<0.01;***p<0.001*.

Growth conditions had no significant effect on calculated water loss during transition from HL to LL (excessive transpiration, ΔW) or calculated CO_2_ loss during transition from LL to HL (forgone photosynthesis, ΔC) (Tables 1 and S3). ⊗W and ⊗C correlated with ct_1/2_ and ot_1/2_, respectively (Table 2). ΔW correlated positively with *a_op_* (r^2^ = 0.397, Figure 4d) and negatively with A_sat_, while ΔC correlated positively with g_sat_ and negatively with SD (Table 2). During light transitions, the C_3_ grass tended to lose more water under all treatments and fix less CO_2_ under high light relative to the C_4_ grasses (Table S3).

### Guard cell K^+^ flux was affected similarly by gCO_2_ but differently by LL in C_3_ and C_4_ grasses

*P. bisulcatum* (C*_3_*) maintained a higher net K^+^ influx rate into guard cells compared to the three C_4_ grasses under HL conditions (Figure 5a, Tables 1 and S3). Relative to the control treatment, net K^+^ influx rate was higher under gCO_2_ for all four species and lower under LL only in *P. bisulcatum* (Figure 5a, Tables 1 and S3). Net K^+^ influx positively correlated with g_s_ (r^2^ = 0.563, Figure 5b), *a_op_* (r^2^ = 0.830, Figure 5b), g_max_ (r^2^ = 0.669), ΔW (r^2^ = 0.588), and OD (r^2^ = 0.365) (Table 2).

**Fig. 5.**
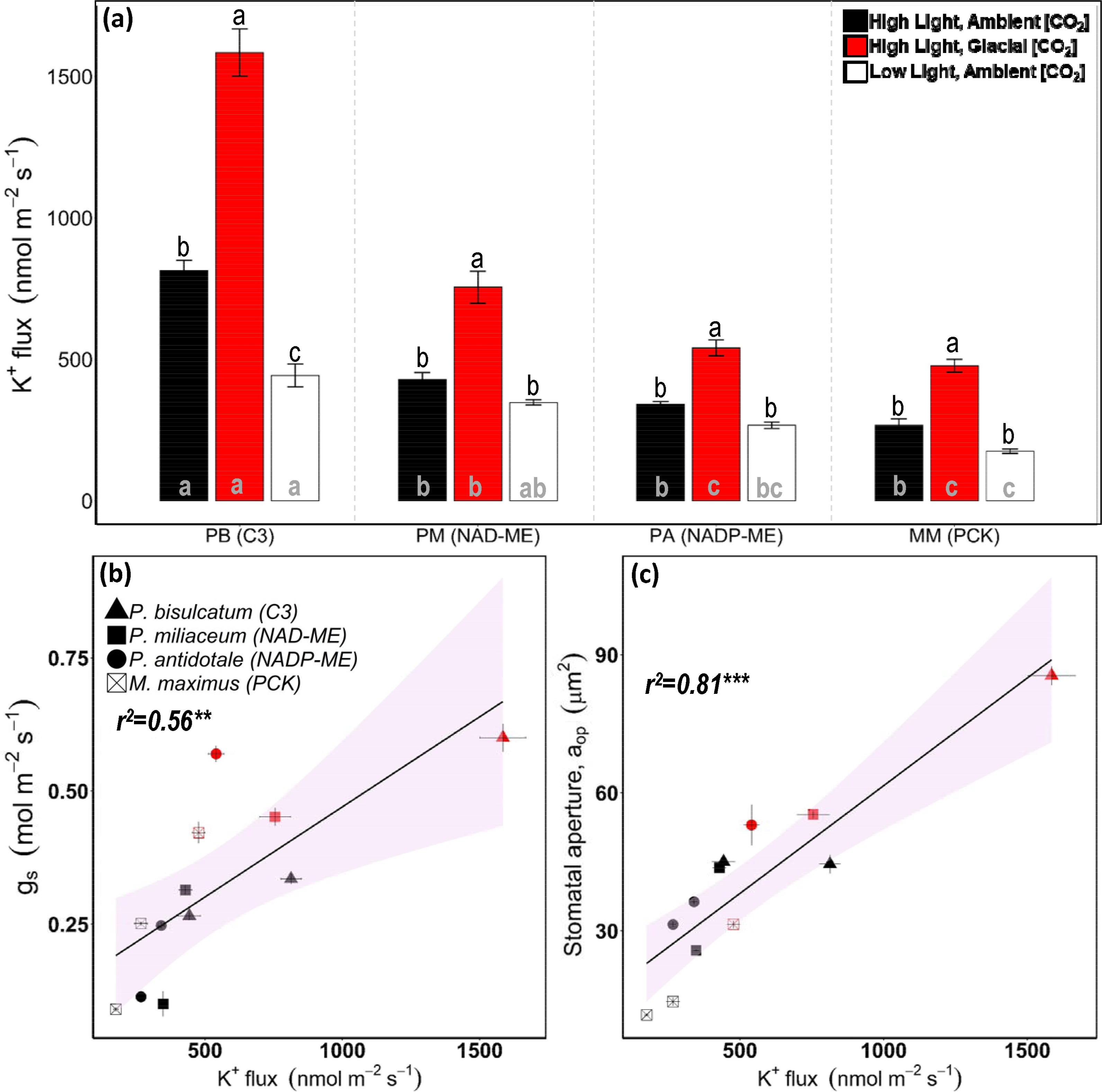
Net K^+^ influx of guard cells (**a**) and its relationship with stomatal conductance (**b**) and stomatal aperture (**c**) in four grass species acclimated to glacial [CO_2_] or low light. Values are means ± SE of n=7-9 stomata and ranked using Tukey’s HSD *post hoc* test at *p* ≤ 0.05. Fitted linear models are as follows: (**b**) g_s_ = 0.0003 × *K^+^flux* + 0.1319, and (**c**) *a _op_* = 0.0494 × *K^+^flux* + 12.87. Other details are described for Fig^X^ure 4.

### Determinants of iWUE across C_3_ and C_4_ grasses

Overall, *iWUE* showed strong negative correlations with g_s_ (r^2^ = 0.398, Figure 6b), *a_op_* (r^2^ = 0. 472, Figure 6c) and guard cell K^+^ influx (r^2^ = 0.587, Figure 6d), and weakly correlated with OD (+ve, r^2^ = 0.213) and A_net_ (-ve, r^2^ = 0.179, Figure 6a) (Table 2). When the key stomatal traits were analysed by PCA, *iWUE* and g_s_ were diametrically opposite (Figure 6e). The parameters *a_op_*, g_s_, g_max_ and K^+^ influx clustered together, while *a_max_*, g_sat_ and SI clustered together. The first (*a_op_*) cluster was flanked by SS/ot_1/2_ and SD/ct_1/2_. Thus, gCO_2_ had an opposite and stronger influence on stomatal traits than LL in the tested grass species (Figure 6e).

**Fig. 6.**
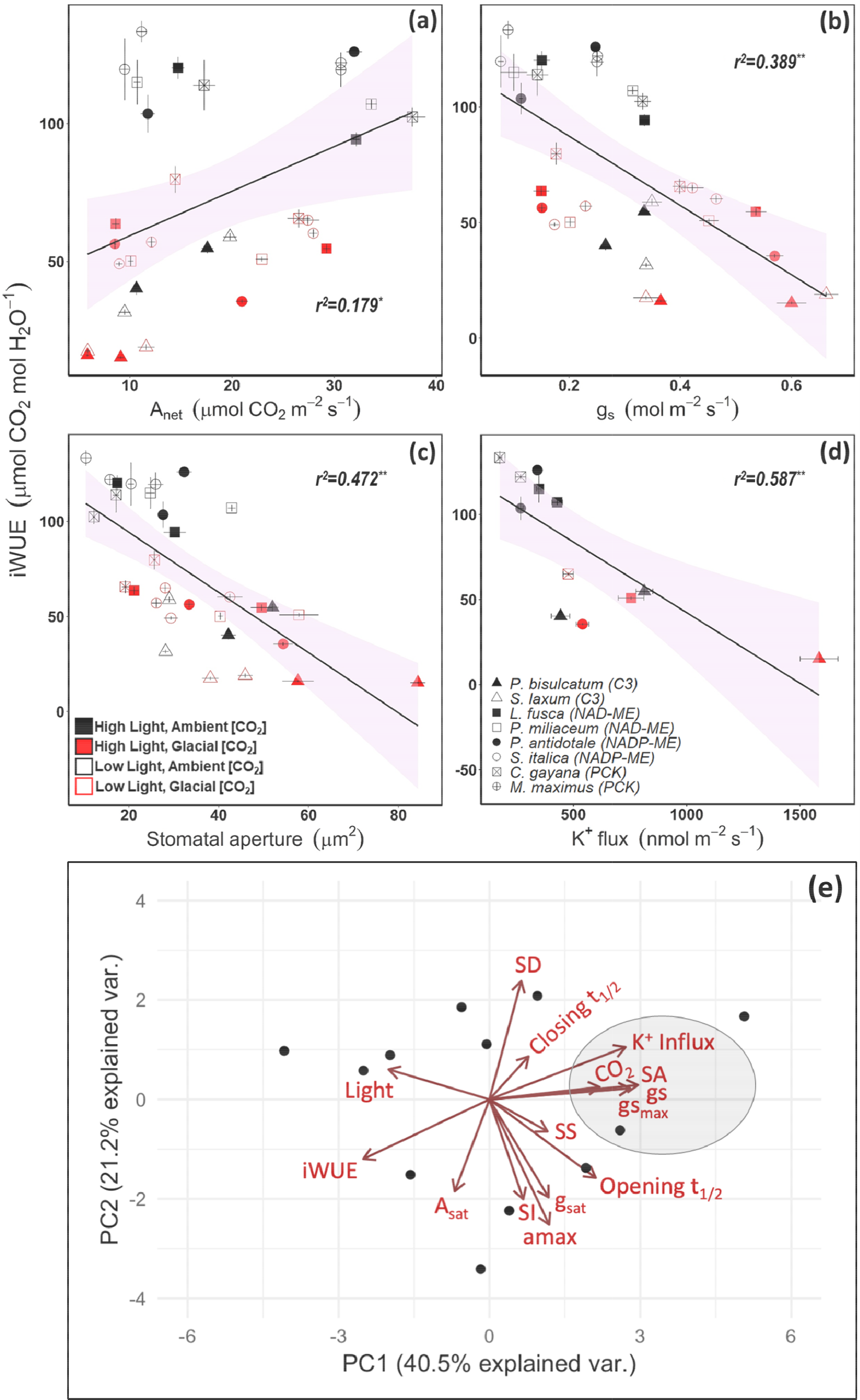
Relationships of intrinsic water-use efficiency (*iWUE*) with (**a**) net photosynthetic rates, (**b**) stomatal conductance, (**c**) stomatal aperture, and (**d**) net K^+^ guard cell influx in eight grasses grown at glacial [CO_2_] or low light. Points represent species means at each growth condition ± SE and shaded regions represent the 95% confidence interval of the linear models. Fitted linear model equations were as follows: (**a**) *iWUE* = 1.6167 × *A_net_* + 43.216; (**b**) *iWUE* = 117.36 – 149.8 ×; (**c**) *iWUE* = 126.04 – 1.5842 × *a_op_*; and (**d**) *iWUE* = 125.27 – 0.0829 × *K^+^flux*. *\*p<0.05; **p<0.01; ***p<0.001*. (**e**) PCA biplot of stomatal traits underpinning *iWUE* was generated using *pca.comp* and visualised using *ggbiplot* in R. The influence of key traits on the principle components separating each sample, across the two principle components accounting for the largest variation. Arrows show the direction and strength of the influence of each trait, while the black circles show where each sample falls across the two principle components.

## DISCUSSION

### Determinants of grass stomatal conductance and responses

This study used eight closely-related C_3_ and C_4_ grasses to explore stomatal characteristics and acclimation to low [CO_2_] and low light, and how these affect stomatal conductance (g_s_) and intrinsic leaf water use efficiency (*iWUE*). C_4_ grasses tended to have lower g_s_, operational stomatal aperture (*a_op_*) and guard cell K^+^ influx rate, and higher CO_2_ assimilation rates (A_net_) and *iWUE* relative to C_3_ grasses. In addition, g_s_ was equally sensitive to gCO_2_ among the grasses, but was more sensitive to LL in C_4_ than C_3_ species. Stomatal size (SS) and stomatal density (SD) showed no trend between the two photosynthetic types. These results confirm what was previously reported in a small subset of species possessing dumbbell-shaped stomata (Maherali *et al*., 2002; Malone *et al*., 1993).

More significant than the differences in these species were the overall correlations and how they varied with the growth environment. For all species, g_s_ depended most on *a_op_*, density of open stomata (OD) and stomatal opening half-time (ot_1/2_). In turn, *a_op_* was largely predicted by the guard cell K^+^ influx rate (r^2^ = 0.830, Figure 5b), and correlated with ot_1/2_ and SS. Although C_4_ grasses generally operated with lower g_s_ and *a_op_* at current aCO_2_, they showed a greater potential to open stomata (g_sat_) relative to maximal stomatal conductance (g_max_), indicating heightened stomatal sensitivity and control as a result of the C_4_-CCM. Finally, species with more stomata (higher SD) had smaller stomata (lower SS) and opened their stomata faster on transition to HL (lower ot_1/2_).

### Stomatal aperture and guard cell K^+^ influx

During stomatal opening and closing, K^+^ fluxes across the guard cell membranes are key determinants of the complex signalling pathways in response to variations of CO_2_, light and other stimuli (Kim *et al*., 2010). Hence, the strong correlations of *a_op_* and g_s_ measured under growth conditions with guard cell K^+^ influx (Table 2, Figure 5) are based on well-understood guard cell function, whereby K^+^, anion and solute influx are accompanied by H_2_O uptake leading to stomatal opening (Chen *et al*., 2017; Chen *et al*., 2012). What is particularly significant in our study is that g_s_ measured *in planta* using leaf gas exchange correlated well with K^+^ influx measured on epidermal peels (Figure 5). This indicates that K^+^ homeostasis of guard cells of isolated epidermal peels generally reflects *in vivo* leaf gas exchange. This was true for the C_3_ grass under all conditions, and for C_4_ grasses under control and gCO_2_, but not LL, conditions.

Stomata have distinct responses and signalling pathways for CO_2_ and light (Assmann and Jegla, 2016; Engineer *et al*., 2016). In our study, higher guard cell K^+^ influx at gCO_2_ relative to aCO_2_ was proportional to g_s_, indicating a common gCO_2_ effect towards increased guard cell K^+^ uptake and signalling response in grasses. However, there was different stomatal acclimation and possibly signalling responses to light between C_3_ and C_4_ grasses because K^+^ influx was not affected by growth at LL in C_4_ grasses while it decreased in proportion with g_s_ in the C_3_ grass (Figure 5). These results may have useful implications for future investigation on the way stomata respond and acclimate to CO_2_ and light in C_3_ and C_4_ grasses.

In addition to different signalling mechanisms, possible explanations for the differential stomatal responses to [CO_2_] relative to light among the grass species may relate to guard cell wall properties. In vascular plants, stomatal cell walls undergo reversible deformation during opening and closing, requiring a strong and flexible structure. The composition (e.g., cellulose, pectin, lignin, phenolics) of guard cell wall differs between phylogenetic groups impacting stomatal function (Rui *et al*., 2018). Noteworthy, whilst C_3_ wheat and C_4_ sorghum possess similar type III dumbbell shaped stomata, wheat guard cells exhibited different relative retardance (cellulose crystallinity) than sorghum (Shtein *et al*., 2017). The relevance of these differences to other C_3_ and C_4_ grass species is yet to be established. Moreover, polar stiffening is emerging as a greater player in guard cell movement than radial stiffening (Carter *et al*., 2017). Understanding how signalling pathways and cell wall composition of guard cells differ between C_3_ and C_4_ grasses and acclimate to the environment may potentially be exploited for crop improvement and will be the subject of future research.

### Stomatal opening speed

Stomata constantly respond to changing light intensity on a scale of seconds through to seasons. Stomatal opening and closing usually occur at a slower rate than photosynthetic activation (Henry *et al*., 2020), creating asynchrony between CO_2_ uptake and H_2_O loss which compromises *iWUE*. Hence, dynamic stomatal responses are critical for the optimisation of *iWUE*, in addition to advanced stomatal morphology and patterning (Cai *et al*., 2017; Lawson and Vialet-Chabrand, 2019; Slattery *et al*., 2018; Vialet-Chabrand *et al*., 2017). In this study, leaves with more open stomata (higher *a_op_*) tended to have lower *iWUE*, slower opening (i.e., greater ot_1/2_), larger excessive transpiration and forgone photosynthesis on transition to HL (Tables 2, S2 and S3). Under gCO_2_ and LL, stomatal opening was generally faster (i.e., smaller ot_1/2_) in CO_2_-saturated C_4_ grasses relative to their C_3_ counterparts (Figure 4b). In contrast, CO_2_-limited C_3_ leaves showed slower stomatal opening at gCO_2_, possibly due to hydraulic constraints as highlighted below.

C_4_ photosynthesis is more complex than C_3_ photosynthesis, involving both C_3_ and C_4_ metabolic cycles operating across two photosynthetic tissues (Hatch, 1987). Hence, efficient C_4_ photosynthesis requires the establishment of substantial metabolite gradients (Arrivault *et al*., 2016; Leegood and von Caemmerer, 1988). Accordingly, it has been shown that C_4_ plants generally have less photosynthetic efficiency and productivity than C_3_ plants under dynamic irradiance, such as sun flecks (Kubásek *et al*., 2013). However, a thorough examination, revealed that C_4_ photosynthesis is buffered against light transients due to the flexible use of the C_4_ acid decarboxylases (Bellasio and Griffiths, 2013; Wingler *et al*., 1999) and reducing equivalents stored in the large metabolite gradients (Slattery *et al*., 2018). Our results concur with the latter consideration, and further suggest that C_4_ grasses can efficiently utilise high light transients due to their highly responsive stomata. Amongst the three subtypes of C_4_ grasses, the PCK species showed faster stomatal opening in response to HL. The PCK subtype is characterised by high photosynthetic efficiency at LL (Sagun *et al*., 2019; Sonawane *et al*., 2018) and more efficient Rubisco (Sharwood *et al*., 2016). Moreover, the two PCK species had smaller stomatal size, which is in line with the general view that the rapidity of g_s_ responses in dumbbell-shaped guard cells in grasses could be attributed to size (McAusland *et al*., 2016).

### Stomatal closing speed

Stomatal ct_1/2_ showed no consistent relationship with measured parameters regardless of photosynthetic type or growth conditions. Rather, C_3_ and NAD-ME grasses had more excessive transpiration (ΔW) due to their slower stomatal closure rate on transition to LL. This may highlight the CO_2_ limitation of the C_3_ *versus* C_4_ pathway and of NAD-ME *versus* NADP-ME and PCK subtypes (Pinto *et al*., 2016; Sonawane *et al*., 2018). Slower stomatal closing rate may also be related to hydraulic conductivity (Taylor *et al*., 2018). Although stomatal ct_1/2_ did not directly correlate with any parameter (other than ot_1/2_), there were some interesting observations. Firstly, ΔW derived from ct_1/2_ and transpiration rates at initial and final steady states, correlated strongly with guard cell K^+^ influx rate (r^2^ = 0.588). Secondly, grass leaves with more frequent stomata (high SD) showed less forgone photosynthesis (ΔC) on opening to HL (r^2^ = 0.292). Among C_3_ kidney-shaped stomata, ΔW and stomatal closing were not correlated (Deans *et al*., 2019). These aspects warrant further investigation using a larger set of C_3_ and C_4_ species.

### Trade-off between SS and SD

So far, we argued that g_s_ responses were dominated by *a_op_* and underpinned by guard cell K^+^ fluxes. Moreover, *a_op_* correlated with SS (Figure 3b). Smaller SS is a well-characterized strategy to speed up solute flux in and out of guard cells by increasing the surface area to volume ratio (Chen *et al*., 2012), and shown to occur in several species (Lawson and Blatt, 2014; Papanatsiou *et al*., 2016; Tanaka *et al*., 2013). However, SS exhibited no significant correlations with g_s_, g_sat_ or g_max_ in our study (Table 2). Hence, the advantage of grass stomata structure may enable small changes in guard cell width to translate into larger changes in *a_op_* (Hetherington & Woodward, 2003; Franks & Farquhar, 2007). This strategy enables grass stomata to rapidly respond under gCO_2_.

In line with other studies (Franks and Beerling, 2009a; Franks *et al*., 2009; Franks *et al*., 2012), SS negatively correlated with SD in our study (Figure 3c). Notably, the two PCK species tended to have smaller SS and higher SD relative to the other C_4_ grasses (Figure 2). The particular interest in SD stems from its apparent environmental plasticity and the prospect that genetic manipulation of SD may impact crop *iWUE* (Buckley *et al*., 2020). Indeed, genetically manipulated C_3_ plants such as rice and wheat with moderate SD reductions were shown to conserve water without suffering a yield penalty (Caine *et al*., 2019; Dunn *et al*., 2019). In the current study, SD was generally not affected by gCO_2_, and could not predict g_s_. Only when we accounted for the frequency of open stomata, OD, a good relationship was obtained between OD and g_s_ (Table 2). These results agree with the conclusions made by (Dow *et al*., 2014a) that g_s_ response to [CO_2_] is a pore-specific property that is scaled by SD. In our study, OD was a more physiologically relevant attribute than SD.

It has been predicted that g_max_ adapts to long term changes in atmospheric [CO_2_] by adjustments in SS and/or SD. In particular, growth at elevated [CO_2_] reduces photosynthetic capacity in C_3_ plants, and it is suggested that optimisation of operational g_s_ requires concomitant reduction in g_max_ to keep it within the sensitive range of guard cell osmotic pressure (Franks *et al*., 2012). However, our data do not support the predictions about SS and SD adjustments as they both changed little under gCO_2_ or LL, even in C_3_ species (Figure 2). Our results agree with other studies using C_4_ grasses grown at low [CO_2_] (Maherali *et al*., 2002), as well as a large survey of published studies at elevated [CO_2_] where SD varied by an average of 5% whilst g_s_ declined by 22% (Ainsworth and Rogers, 2007). Therefore, the global inverse relationship between SD and atmospheric [CO_2_] documented in the fossil records may partly be due to the larger changes in [CO_2_] across the geological timescale (Franks and Beerling, 2009b; Woodward, 1987; Woodward and Kelly, 1995), or to variations among species (Aasamaa and Aphalo, 2016; Evans-Fitz Gerald *et al*., 2016), leaf positions or sun/shade aspects (Poole *et al*., 1996).

### Maximal stomatal conductance

In this study, we estimated maximal stomatal conductance anatomically (g_max_) by measuring SD and *a_max_*, and physiologically (g_sat_) by measuring leaf gas exchange at low CO_2_ and high light (Figure 1). g_sat_ showed no correlation with any measured stomatal traits, while g_max_ weakly correlated with g_s_, SD and ot_1/2_ (Table 2). These results partially fit with work using Arabidopsis SD mutants demonstrating that operational stomatal conductance can be estimated as a function of g_max_, independently of SD (Dow and Bergmann, 2014; Dow *et al*., 2014a; Dow *et al*., 2014b). For the two C_3_ species, g_s_ was a constant fraction of g_sat_ and g_max_ when measured at aCO_2_, while g_s_/g_sat_ was higher at gCO_2_ (Figures 1f-h, Tables 1 and S1). For the C_4_ species, the g_s_/g_sat_ and g_s_/g_max_ ratios were higher at gCO_2_ and lower at LL. Moreover, the C_3_ species had higher g_s_/g_sat_ while the C_4_ species had higher g_sat_/g_max_ (Figures 1f-h, Tables 1 and S1). These results indicate that while C_3_ grasses operate with higher g_s_, the C_4_ grasses have a greater potential (measured as g_sat_) for stomatal opening relative to their g_max_. In turn, these results indicate the greater sensitivity of C_4_ stomata to open at gCO_2_ which reflects both their low CO_2_-limitation due to the CCM and their reduced risk of hydraulic failure at low CO_2_. Both factors allow C_4_ stomata to open more in response to gCO_2_ (Pinto *et al*., 2014). These conclusions are supported by previous work showing that g_s_ was more sensitive to changing intercellular CO_2_ in C_4_ relative to C_3_ species within the dicot genus *Flaveria* (Huxman and Monson, 2003; Vogan and Sage, 2011).

### Determinants of grass iWUE

Across all species and treatments, *iWUE* was most affected by g_s_, *a_op_* and guard cell K^+^ influx, which all strongly correlated among each other. Overall, *iWUE* correlated more strongly with g_s_ than A_net_. The correlation with A_net_ here was stronger than what has been previously reported in other studies utilising C_4_ grasses (Cano *et al*., 2019). C_4_ photosynthesis is largely CO_2_-saturated at aCO_2_, hence changes in g_s_ have little effect on A_net_, but large effects on *iWUE* (Ghannoum, 2016; Li *et al*., 2017). In contrast, *iWUE* is strongly affected by both A_net_ and g_s_ in C_3_ plants (Farquhar and Sharkey, 1982). The inclusion of two C_3_ grasses and the gCO_2_ treatment has strengthened the influence of A_net_ on *iWUE* in our set of mostly C_4_ grasses. Consequently, *iWUE* increased as A_net_ increased and g_s_, *a_op_*, SS and K^+^ flux decreased (Figure 6).

## Conclusions

In conclusion, we reveal new insight on the variation in stomatal behaviour of C_3_ and C_4_ grasses, highlighting both conserved and distinct responses determined by photosynthetic type (C_3_ and C_4_) and environmental condition (gCO_2_ and LL). Across species and treatments, stomatal conductance (g_s_) and *iWUE* correlated well with guard cell K^+^ influx and *a_op_*, but poorly with SD. Grasses with smaller stomata had smaller *a_op_* and faster stomatal opening on transition to HL, which allowed them to maintain greater steady-state and dynamic *iWUE*. In addition, C_4_ grasses exhibited different responses of g_s_ and guard cell K^+^ influx to LL compared to C_3_ stomata allowing them to maintain higher *iWUE* at LL. These mechanistic links between stomatal traits highlight future avenues for improving photosynthesis and water use efficiency via improved understanding of stomatal function in C_3_ and C_4_ cereal species.

## AUTHOR CONTRIBUTION

All authors contributed to experimental design. WKI and AJWL carried out the experimental work and data analysis. WKI and OG wrote the manuscript with contribution from AJWL and ZHC.

## ACKNOWLEDGEMENTS

This research was funded by the Australian Research Council through the Centre of Excellence for Translational Photosynthesis (CE1401000015) awarded to OG and DECRA award to ZHC (DE1401011143). WKI was supported by a PhD scholarship jointly awarded by the Centre of Excellence for Translational Photosynthesis and the Hawkesbury Institute for the Environment, Western Sydney University (WSU). We gratefully acknowledge the technical support of Fiona Koller, Dr Chenchen Zhao, Dr Shengguan Cai, Abigail Tuddao and Mark Christophere Tarculas at WSU.

